# CRISPR-Cas off-target detection using Oxford Nanopore sequencing - is the mitochondrial genome more vulnerable to off-targets?

**DOI:** 10.1101/741322

**Authors:** Sandeep Chakraborty

## Abstract

Oxford Nanopore sequencing of DNA molecules is fast gaining popularity for generating longer reads, albeit with higher error rates, in much lesser time, and without the error introduced by PCR-amplification. Recently, CRISPR-Cas9 has been used to enrich genomic regions (nCATS [1]). This was applied on 10 genomic loci (median length=18kb). Here, using the sequencing data (Accid:PRJNA531320), it is shown that the same flow can be used to identify CRISPR-Cas9 off-target edits (OTE). OTEs are an important, but unfortunately underestimated, aspect of CRISPR-Cas gene-editing. An OTE in the mitochondrial genome is shown having 7 mismatches with one of the 10 gRNAs used (GPX1), having as much enrichment as the targeted genomic loci in some samples. Previous study has shown that Cas9 bind to off-targets having as many as 10 mismatches in the PAM-distal region. This OTE has not been reported in the original study (still a pre-print), which states that sequences from parts other than the target locations arise ‘from ligation of nanopore adaptors to random breakage points, with no clear evidence of off-target cleavage by Cas9’ [1], Furthermore, a lot of reads aligning to the mitochondrial genome (sometimes full length) are inverted after the edit. It remains to be seen if these are bona fide translocations after the Cas9 edit, or ONP sequencing artifacts. This also raises the question whether the mitochondrial genome is more prone to off-targets by virtue of being non-nuclear. Another locus in ChrX (13121412) has only 1 mismatch with the second BRAF gRNA (GACCAAGGATTTCGTGGTGA). Although the number of reads for this OTE is less, its very unlikely this is random since it happens 8 out of 11 samples. With the increasing use of (TALEN/ZFN/CRISPR-Cas9) on human subjects, this provides a fast method to quickly query gRNAs for off-targets in cells obtained from the patient, which will have their own unique off-targets due to single nucleotide polymorphism or other variants.

## Introduction

Fluctuations in electrical conductivity generated as DNA strands pass through a biological pore, like a protein embedded in a membrane has been commercialized by Oxford Nanopore (ONP) (Nanopore sequencing - minION flow cell) [2]. ONP sequencing is gaining popularity due to the fact that it quickly generates long reads, preserving nucleotide modifications, albeit at higher error rates [3]. Due to higher error rates, enrichment is critical to have proper consensus. Recently, CRISPR-Cas9 [4–6] has been used to enrich genomic regions (nCATS [1]. In this method all DNA ends are first dephosphorylated, then the Cas9/guide RNA ribonucleoprotein complex (RNP) is added resulting in DSBs, thereby introducing an DNA end with a 5-phosphate group, wherein nanopore sequencing adaptors are ligated, and then sequenced. Note, that PCR amplification is not needed, thus preserving nucleotide modifications. This was applied on 10 genomic loci (median length=18kb), and the sequencing data were submitted (Accessionid:PRJNA531320).

Off-targets edits (OTE) are an important aspect of CRISPR-Cas gene-editing [7–10]. There are several techniques for the unbiased detection of CRISRP-Cas9 off-targets [11–13]. The presence of the ONP adapters in the above study enable the identification of OTEs, aided by the existence of 11 such “repeated experiments”.

As an example, an OTE in the mitochondrial genome is provided which has the ONP adapter, and has 7 mismatches with one of the 10 gRNAs used. Insertions or deletions between target DNA and gRNA also constitute valid off-targets [14], a feature that is included in *in silico* off-target predictors [15]. Another OTE is ChrX for the BRAF gRNA with only one mismatch. While it stated that sequences from parts other than the target locations arise ‘from ligation of nanopore adaptors to random breakage points, with no clear evidence of off-target cleavage by Cas9’ [1], it is very unlikely that all samples will have the breakage at the same point.

With the increasing use of (TALEN/ZFN/CRISPR-Cas9) on human subjects, this provides a fast method to quickly query gRNAs for off-targets in cells obtained from the patient, which will have their own unique off-targets due to single nucleotide polymorphism or other variants.

## Results and discussion

### Reads matching to the 10 edited-genomic loci

Previous work [1] used 2 gRNAs for each of 10 genomic loci (Table 1). The reads matching to one such loci (SLC12A4) with high significance were analyzed (Fig 1). This corroborated two things:

1. The error rare is *>* 10% - since most alignments were about 90% or less.
2. The Cas9 mostly edits about 2 bases to the left of the PAM site.

**Table 1:** **Ten gRNAs used in previous study [1]:** The sequences within these coordinates are in SI:TARGET.fa

| Gene/Chr | Accid | start | end | size |
| --- | --- | --- | --- | --- |
| GSTP1CHR11 | GI.568815271.REF.NT_167190.2. | 13051355 | 13069174 | 17819 |
| KRT19CHR17 | GI.568802181.REF.NT_010783.16. | 14581543 | 14599732 | 18189 |
| TPM2CHR9 | GI.568815299.REF.NT_008413.19. | 35667526 | 35687167 | 19641 |
| GPX1CHR3 | GI.568815345.REF.NT_022517.19. | 49342526 | 49356170 | 13644 |
| SLC12A4CHR16 | GI.568802190.REF.NT_010498.16. | 21571688 | 21596077 | 24389 |
| CHR5DELETION | GI.568815317.REF.NT_034772.7. | 8268286 | 8286975 | 18689 |
| CHR7DELETION | GI.806527423.REF.NW_012132919.1. | 170963 | 92107200 | 91936237 |
| BRAFCHR7 | GI.568815306.REF.NT_007933.16. | 78268436 | 78280774 | 12338 |
| KRASCHR12 | GI.568815268.REF.NT_009714.18. | 18153329 | 18170070 | 16741 |
| TP53CHR17 | GI.568802188.REF.NT_010718.17. | 7177042 | 7193140 | 16098 |

**Figure 1:**
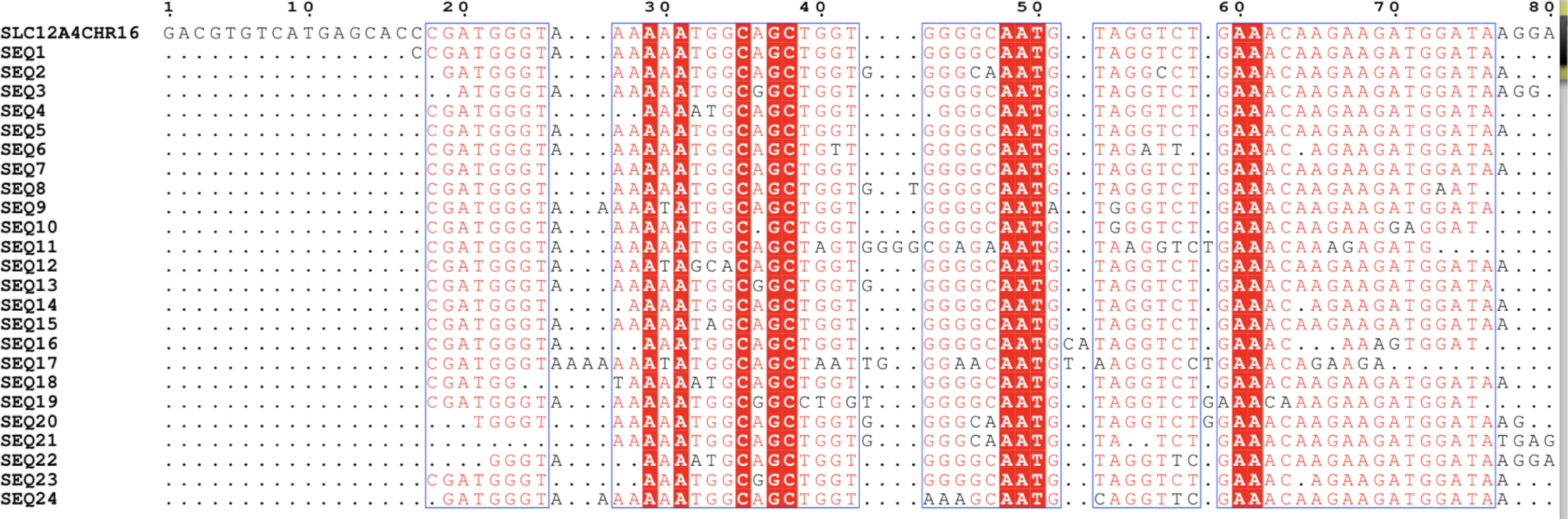
**An estimate of the error rates in ONP:** Most reads match to the reference genome with about 90% homology, giving an error rate of about 10%. This small fragment from the SLC12A4 gene on CHR16 demonstrates that the consensus clearly matches to the reference, inspite of the error rate. This also shows that the Cas9 mostly edits about 2 bases to the left of the PAM site.

The ONP adapter was prefixed to the Cas9 edit site, deduced to be TCAGTATTGCTTCGTTCAGT-TAC. Next, reads that matched to the targeted genomic loci (SI:TARGET.fa) were ignored. The remaining reads were then selected only if they had the ONP adapter, and aligned to the human genome.

### Reads matching to the mitochondrial genome - off-target common to many samples

The alignment revealed that a significant number of reads matched to the mitochondrial genome with high significance in many cell lines (Table 2). Interestingly, many of these reads started around loci 9639, which has 7 mismatches (Fig 2) with one of the used gRNAs (Table tableinfo,GPX1). Only 2 of the 7 mismatches are located in the 12bp proximal to the PAM. Note, previous work shows that Cas9 can bind ‘*with as many as nine consecutive mismatches in the sgRNA guiding*’ [9]. Also, Cas9 cleaves for a wide range of PAMs [16], including the NGA PAM in this case. In fact, for one gRNA (gP), *in vitro* studies showed that Cas9 was targeting several sites with as many as seven mismatches [16]. Also, a lot of reads aligning to the mitochondrial genome (sometimes full length) are inverted after the edit. It remains to be seen if these are bona fide translocations after the Cas9 edit, or ONP sequencing artifacts. The reads aligning to the mitochondrial genome can be found at SI:ALLMITO.fa.

**Table 2:** **ONP sequencing data from 11 samples:** The data is from Accid:PRJNA531320. Number of reads matching to a targeted loci (GPX1) of length 13644bp is compared to reads that match with the mito-chondrial genome (MITOGENO,16570bp). The cutoff values is set to Blastscore=1000 to choose reasonably large reads. There are significant reads which start at 9639 in the MITOGENO, which has 7 mismatches with one of the gRNAs used to edit the GPX1 loci, indicating that these are Cas9 off-targets. Another locus in ChrX (13121412) has only 1 mismatch with the second BRAF gRNA (GACCAAGGATTTCGTGGTGA). These reads all have the ONP adaptor. epithelial fibrocystic disease=EPIFIB, epithelial adenocarcinoma=EPIADE, caucasian, mammary gland=CMG

| Accid | Description | Total reads | SLC12A4 | GPX1 | Mito | 9639 loci | ChrX |
| --- | --- | --- | --- | --- | --- | --- | --- |
| SRR8863250 | GM12878 BLOODlymphocyte | 188570 | 0 | 0 | 498 | 0 | 0 |
| SRR9167335 | GM12878 BLOODlymphocyte | 14522 | 55 | 76 | 32 | 0 | 0 |
| SRR9167337 | GM12878 BLOODlymphocyte | 188570 | 0 | 0 | 507 | 0 | 0 |
| SRR9167338 | GM12878 BLOODlymphocyte | 89433 | 245 | 160 | 339 | 17 | 1 |
| SRR9167340 | MCF-7 69y EPIADE CMG | 84201 | 406 | 122 | 563 | 22 | 1 |
| SRR8863251 | GM12878 BLOODlymphocyte | 89432 | 239 | 156 | 333 | 43 | 1 |
| SRR9167339 | MDA-MB-231 51y1 EPIADE CMG | 236363 | 536 | 488 | 1943 | 62 | 2 |
| SRR8863247 | MCF-7 69y EPIADE CMG | 84198 | 398 | 119 | 552 | 100 | 2 |
| SRR8863249 | MDA-MB-231 51y EPIADE CMG | 229208 | 503 | 455 | 1829 | 145 | 2 |
| SRR9167336 | MCF-10A 36y A EPIFIBe CMG | 70542 | 740 | 754 | 3912 | 168 | 4 |
| SRR8863248 | MCF-10A 36y EPIFIBe CMG | 70542 | 703 | 729 | 3819 | 645 | 5 |

**Figure 2:**
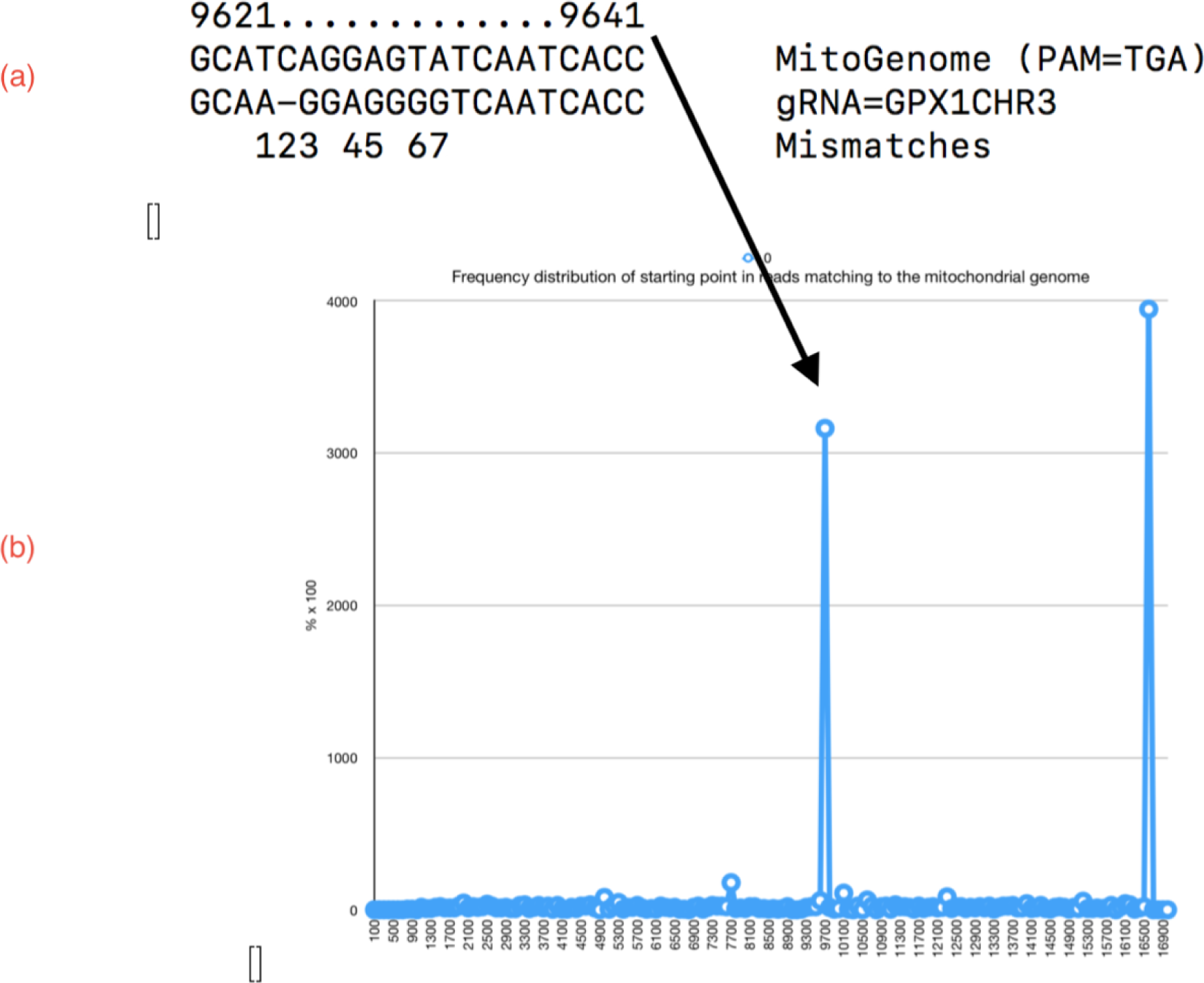
**Off-target in the mitochondrial genome: (a):** The genomic loci has 7 mismatches with one of the gRNAs (GPX1 on chr3), and only 2 in the 12bp proximal to the PAM. Note, previous work shows that Cas9 can bind ‘*with as many as nine consecutive mismatches in the sgRNA guiding*’ [9]. Also, Cas9 cleaves for a wide range of PAMs [16], including the NGA PAM in this case. **(b):** Frequency distrubtion of the start of the reads with respect to the mitochondrial genome (16570bp), reveals a significant number of reads begin around 9639. These reads also have the ONP adapter pre-fixed to them.

### Off-target on the X chromosome

Some of the gRNAs have OTEs with low mismatches. In fact, it would be surprising not to find OTEs for such gRNA. As an example, BRAF (gRNA=GACCAAGGATTTCGTGGTGA) on chr7 has 19 nt exact match to the X-chromosome at 13121412. 8 out of 11 samples have reads matching to this coordinate. This cannot be random (Table 2). The read ids are in SI:OTEChrX.list.

## Conclusion

The method (nCATS) suggested recently [1] to enrich genomic regions of interest also provides an easy method to quickly query off-target edits. With the increasing use of (TALEN/ZFN/CRISPR-Cas9) on human subjects, this provides a fast method to quickly query gRNAs for off-targets in cells obtained from the patient, which will have their own unique off-targets due to single nucleotide polymorphism [17] and other variants.

## Materials and methods

The sequencing data (11 samples) were downloaded from NCBI (PRJNA531320). The genomic coordinates encapsulated between the 2 gRNAs for each of 10 genomic loci were extracted (SI:TARGET.fa). In order to search for off-targets, reads matching to the targeted genomic coordinates were first discarded. Next, reads were clustered based on their homology to each other - this was to look for enriched loci, and avoid random breaks (since off-targets are being identified). The mitochondrial genome was obtained from Accid:GI.251831106.REF.NC 012920.1.(16570bp). MSA figures were generated using the ENDscript server [18].

## Supporting information

SI Data - sequences

## References

1. Gilpatrick T, Lee I, Graham JE, Raimondeau E, Bowen R, et al. (2019) Targeted nanopore sequencing with cas9 for studies of methylation, structural variants and mutations. BioRxiv : 604173.

2. Jain M, Olsen HE, Paten B, Akeson M (2016) The oxford nanoporeminION: delivery of nanopore sequencing to the genomics community. Genome biology 17: 239.

3. Lu H, Giordano F, Ning Z (2016) Oxford nanoporeminION sequencing and genome assembly. Genomics, proteomics & bioinformatics 14: 265–279.

4. Barrangou R, Fremaux C, Deveau H, Richards M, Boyaval P, et al. (2007) CRISPR provides acquired resistance against viruses in prokaryotes. Science 315: 1709–1712.

5. Jinek M, Chylinski K, Fonfara I, Hauer M, Doudna JA, et al. (2012) A programmable dual-RNA–guided DNA endonuclease in adaptive bacterial immunity. Science 337: 816–821.

6. Mali P, Yang L, Esvelt KM, Aach J, Guell M, et al. (2013) RNA-guided human genome engineering via Cas9. Science 339: 823–826.

7. Fu Y, Foden JA, Khayter C, Maeder ML, Reyon D, et al. (2013) High-frequency off-target mutagenesis induced by CRISPR-cas nucleases in human cells. Nature biotechnology 31: 822–826.

8. Pattanayak V, Lin S, Guilinger JP, Ma E, Doudna JA, et al. (2013) High-throughput profiling of off-target DNA cleavage reveals rna-programmed Cas9 nuclease specificity. Nature biotechnology 31: 839.

9. Kuscu C, Arslan S, Singh R, Thorpe J, Adli M (2014) Genome-wide analysis reveals characteristics of off-target sites bound by the Cas9 endonuclease. Nature biotechnology 32: 677.

10. Chakraborty S (2018) Inconclusive studies on possible CRISPR-cas off-targets should moderate expectations about enzymes that have evolved to be non-specific. Journal of Biosciences 43: 225–228.

11. Wienert B, Wyman SK, Richardson CD, Yeh CD, Akcakaya P, et al. (2019) Unbiased detection of CRISPR off-targets *in vivo* using DISCOVER-Seq. Science 364: 286–289.

12. Tsai SQ, Nguyen NT, Malagon-Lopez J, Topkar VV, Aryee MJ, et al. (2017) CIRCLE-seq: a highly sensitive in vitro screen for genome-wide CRISPR-Cas9 nuclease off-targets. Nature Methods 14: 607.

13. Tsai SQ, Zheng Z, Nguyen NT, Liebers M, Topkar VV, et al. (2015) GUIDE-seq enables genome-wide profiling of off-target cleavage by CRISPR-Cas nucleases. Nature biotechnology 33: 187.

14. Lin Y, Cradick TJ, Brown MT, Deshmukh H, Ranjan P, et al. (2014) CRISPR/Cas9 systems have off-target activity with insertions or deletions between target DNA and guide RNA sequences. Nucleic acids research 42: 7473–7485.

15. Bae S, Park J, Kim JS (2014) Cas-offinder: a fast and versatile algorithm that searches for potential off-target sites of Cas9 rna-guided endonucleases. Bioinformatics 30: 1473–1475.

16. Akcakaya P, Bobbin ML, Guo JA, Lopez JM, Clement MK, et al. (2018) In vivo CRISPR-cas gene editing with no detectable genome-wide off-target mutations. bioRxiv : 272724.

17. Lessard S, Francioli L, Alfoldi J, Tardif JC, Ellinor PT, et al. (2017) Human genetic variation alters CRISPR-Cas9 on-and off-targeting specificity at therapeutically implicated loci. Proceedings of the National Academy of Sciences : 201714640.

18. Robert X, Gouet P (2014) Deciphering key features in protein structures with the new endscript server. Nucleic acids research 42: W320–W324.

